# Edi-Set-Flow: detecting differential RNA-editing at scale

**DOI:** 10.64898/2026.11.02.763838

**Authors:** Jacob E. Munro, Melanie Bahlo, Brendan R. E. Ansell

## Abstract

**Background:** Adenosine-to-inosine (A-to-I) RNA editing is a post-transcriptional modification catalysed by ADAR enzymes that can alter codons, splicing patterns, and RNA secondary structures. RNA editing is essential for innate immunity and nervous system development, and its dysregulation is implicated in neurological disorders, cancers, and autoimmune diseases. Despite its biological importance, accurately detecting RNA editing from RNA-seq data remains technically challenging, and robust inference of differential editing across experimental conditions is not straightforward.

**Findings:** We have developed Edi-Set-Flow, a reproducible and scalable pipeline for transcriptome-wide A-to-I RNA editing analysis of bulk RNA-seq data. Implemented in Nextflow, the pipeline takes raw FASTQ files as input, trims reads and filters for quality, aligns reads to the reference genome, and identifies editing sites with JACUSA2. Common genetic variants are excluded using the gnomAD population database. Identified sites are annotated for genetic feature context and predicted functional consequences, and results are summarised in a user-friendly HTML report. The pipeline scales efficiently to hundreds or thousands of samples and is suitable for large datasets such as GTEx.

An accompanying R package enables advanced analyses, including model fitting, hypothesis testing, false discovery rate control, and visualisation, thereby facilitating reliable statistical comparisons of editing across experimental groups. Applying Edi-Set-Flow to GTEx brain RNA-seq data, we uncovered distinct RNA editing signatures across brain tissues and recovered known protein-recoding events.

**Conclusions:** Edi-Set-Flow provides researchers with a robust, user-friendly, scalable, end-to-end solution for efficiently identifying and interpreting biologically meaningful RNA editing events across diverse transcriptomic datasets, for both mouse and human.

## Introduction

Adenosine-to-inosine (A-to-I) RNA editing is a ubiquitous post-transcriptional modification, with millions of editing sites detectable in human RNA sequencing data^1^. A family of enzymes known as adenosine deaminases acting on RNA (ADARs) carries out this process. ADARs can bind to dsRNA and catalyse the chemical conversion of adenosine to inosine, which is recognised by both the cellular translational machinery and short-read sequencing platforms as guanosine^2^. ADARs are present in all metazoan genomes; the human genome encodes three ADAR genes, namely *ADAR* – encoding the ADAR1 p110 (110 kDa) and p150 (150 kDa) isoforms, *ADARB1* – encoding ADAR2, and *ADARB2* – encoding ADAR3, which is not known to be catalytically active but is expressed and can bind dsRNA^3^. The importance of A-to-I editing is underscored by the fact that biallelic loss-of-function variants in both *ADAR* and *ADARB1* result in severe Mendelian disease. Specifically, *ADAR* deficiency causes the autoimmune disorder Aicardi-Goutières syndrome 6 (AGS6, MIM #615010)^4^, and *ADARB1* deficiency causes a developmental and epileptic encephalopathy (MIM #618862)^5^.

The ADAR1 isoforms show a greater preference for editing repetitive elements^6^. In humans, these are primarily *Alu* repeats, which comprise approximately 10% of the genome and are common within introns and 3′ UTRs^7^. Transcription of genes containing pairs of inversely oriented *Alu* repeats can lead to the formation of dsRNA hairpin structures. The primary function of ADAR1 is to prevent these endogenous dsRNAs from inappropriately activating the innate immune response. Cytosolic dsRNA, including that produced during many viral infections, is recognised by pattern recognition receptors such as MDA5 and PKR, leading to the production of type I interferon and induction of antiviral interferon-stimulated genes (ISGs)^8^. ADAR1 limits the recognition by these receptors by editing endogenous dsRNA, thereby acting as a negative regulator of this pathway. Consistent with this role, *Adar1* knockout is embryonically lethal in mice and is associated with severe dysregulation of interferon-mediated innate immunity. In humans, loss of ADAR1 activity in AGS6 results in inappropriate recognition of endogenous dsRNA, chronic type I interferon signalling and increased expression of ISGs. AGS6 is consequently characterised by a neuroinflammatory phenotype that includes intracranial calcification, microcephaly, developmental delay, dystonia, cerebral palsy and seizures, and can phenotypically resemble congenital viral infection^4,9,10^.

ADAR2 (*ADARB1)* displays greater specificity for editing non-repetitive and coding regions^11^. These editing sites are more commonly located within coding regions, where editing can alter the encoded protein sequence through amino acid substitutions. A well-characterised example is the Q/R editing site in *GRIA2*, which encodes a subunit of the AMPA-type glutamate receptor. Editing at this site occurs in almost all transcripts and converts a glutamine residue to arginine, greatly reducing the receptor’s calcium permeability^12^. Loss of *Adar2* in mice causes seizures and premature death, with much of this phenotype attributable to failure to edit the codon corresponding to the *Gria2* Q/R site^7^. In humans, biallelic loss-of-function variants in *ADARB1* cause neurodevelopmental disorder with hypotonia, microcephaly, and seizures (NEDHYMS, MIM #618862)^5^.

A-to-I RNA editing events are detectable in short-read RNA-seq data as A>G substitutions. Either stranded or unstranded RNA-seq data may be used; however, strand-specific library preparation has the advantage that each substitution may be unambiguously assigned to the strand on which it occurred. For unstranded data, the edited strand must be inferred from the transcript model, and in cases of overlapping transcripts in opposite orientations, A>G substitutions from editing cannot be reliably separated from T>C substitutions arising on an overlapping antisense transcript.

Two widely used RNA editing analysis tools exist. The REDItools^13^ suite is widely used to detect RNA-editing sites by applying depth- and frequency-based filtering of A>G counts from BAM alignments, with an optional step to exclude genomic SNV sites using matched WGS. Another popular tool is JACUSA2^14^, which provides a fast, general-purpose tool for detecting RNA base modifications, including A-to-I editing, and enables strand-specific counting of bases supporting a particular substitution from BAM files. JACUSA2 can be used in single-sample modes or in paired RNA-vs-RNA and RNA-vs-DNA modes to detect sites with significantly different levels of RNA editing, and it includes several built-in filters to remove common artefacts that confound site detection in short-read RNA-seq data.

A-to-I editing is typically quantified as the sum of guanosine over the total coverage (sum of guanosine and adenosine) to give an editing rate (also referred to as the ‘frequency of inosine’) at each site per sample. Differential A-to-I editing has been assessed using a range of statistical techniques, including Student’s t-test^15^, Wilcoxon rank-sum test^16^, and Fisher’s exact test^6^. However, because editing rates are estimated from a finite number of reads, both their resolution and precision depend on sequencing depth. This effect is most pronounced at lower depths, where only a small number of editing-rate values are possible, and estimates have greater sampling variability. T-tests and Wilcoxon rank-sum tests are two-group comparison tests applied to per-sample editing rates and do not account for the depth-dependent sampling variability, resulting in reduced statistical power^17,18^. Furthermore, these tests do not allow for additional covariates such as age, sex, or sequencing batch. Approaches based on Fisher’s exact tests instead pool read counts across samples within each group into a single two-by-two contingency table, thereby discarding any between-sample variability. This has been shown to result in inflated false-positive rates that worsen as sequencing coverage increases^18^.

More sophisticated approaches address these limitations by jointly modelling read counts and between-replicate variability within a generalised linear model or mixed model framework. One such example is REDITs^18^, which implements two beta-binomial-based tests: REDIT-LLR, a likelihood-ratio test for two-group comparisons that does not model covariates, and REDIT-Regression, which instead implements the beta-binomial test as a generalised linear model (GLM) incorporating arbitrary fixed-effect covariates (e.g., age, sex). Another tool, CADRES^19^, performs two-condition differential testing using a binomial generalised linear mixed model as implemented in rMATS^20^. CADRES requires paired whole-genome or whole-exome sequencing data along with RNA-seq to identify variant sites that are uniquely present in the RNA data, increasing sequencing requirements and costs. Additionally, the differential modelling does not support fixed-effect covariates.

We sought to fill a gap in the current landscape of differential RNA editing analysis tools: an end-to-end, scalable pipeline for differential RNA-editing analysis that leverages population genetics for pragmatic and cost-effective exclusion of genomic SNPs, allows modelling of more than two experimental groups, can incorporate arbitrary covariates, and effectively controls for false discoveries due to overdispersion in observed editing rates. The pipeline is designed for large-scale analyses involving hundreds to thousands of samples. We have benchmarked a range of GLM-based approaches for detection of differential RNA-editing, and optimised for sensitivity while maintaining adequate control of false discovery rates.

## Methods

### Edi-Set-Flow: Software implementation

Edi-Set-Flow is implemented in Nextflow^21^, a language that enables the execution of reproducible scientific workflows on a wide range of computation backends, including cloud, high-performance computing (HPC) clusters, and local execution. Software containerisation with Docker, Singularity, or Apptainer ensures pipeline results are reproducible across compute environments. On first run, the pipeline automatically retrieves required reference datasets, including the reference genome (supported options: human - hg38, mouse – mm10 and mm39), GENCODE gene annotations, and UCSC repeat tracks. For human analyses only, the pipeline also downloads dbSNP and gnomAD v4.1 variant calls to exclude polymorphic sites, and retrieves the REDIportal atlas of editing-site annotations.

The pipeline takes as input RNA-seq data in either FASTQ format or aligned BAM format, and proceeds through five main stages: 1) Preprocessing, performing trimming and alignment; 2) Discovery, performing detection of candidate sites; 3) Quantification, performing counting of edited and unedited bases; 4) Annotation; and 5) Reporting, which includes optional modelling of site-specific differences in RNA editing between experimental groups (Figure 1). Results are then output in user-friendly HTML and machine-readable formats.

**Figure 1:**
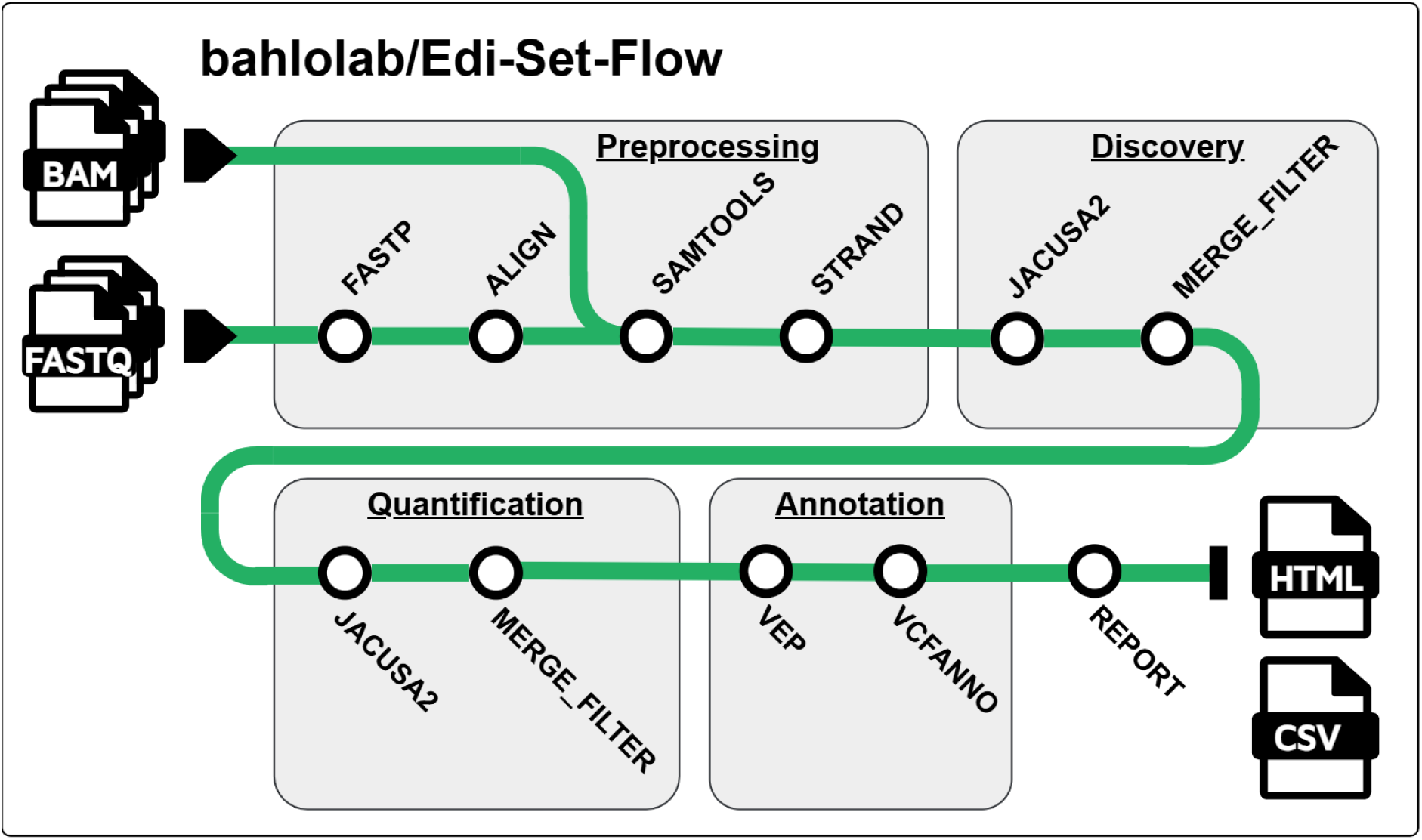
Schematic overview of the Edi-Set-Flow pipeline.

### Edi-Set-Flow: Data preprocessing

Edi-Set-Flow performs optional preprocessing and quality control with fastp (v0.24) ^22^. Users may perform preprocessing in predefined ‘MINIMAL’ or ‘STRICT’ modes. In both modes, reads shorter than 30 bp are discarded, and adapter sequences are trimmed. Additional quality control is applied in STRICT mode: reads are discarded if over 40% of bases are below Phred 15, if there are more than 5 ambiguous bases, and poly-X tails of 7 or more bases are trimmed to remove sequencing artefacts.

Alignment is then performed using either bwa-mem2^23^ (v2.2.1) or STAR^24^ (v2.7.11b, 2-pass mode). STAR is typically the preferred aligner for RNA-seq analyses because it is splice-aware and, when run in “2-pass” mode, can also detect and realign reads at unannotated splice junctions. Following alignment, reads with uncertain mapping quality (MAPQ = 0) are removed, and duplicate, secondary and supplementary read alignments are optionally removed. The alignments are then sorted, and MD tags (indicating base mismatch/deletion) are computed against the reference genome using samtools calmd (v1.21). This last step is required by JACUSA2, which relies on MD tags to identify base mismatches without accessing the reference genome during analysis.

Finally, library strandedness is detected using RSeQC^25^. This step avoids unintentional mis-specification of library strandedness, which can ultimately lead to errors in quantifying A-to-I editing sites by JACUSA2.

### Edi-Set-Flow: Site discovery and quantification

By default, the target genomic regions for site discovery are derived from the union of annotated GENCODE gene features and known REDIportal sites, excluding common single-nucleotide polymorphism sites (MAF > 0.01) from gnomAD v4.1 and dbSNP. Target genomic regions may, however, be otherwise defined by the user. Edi-Set-Flow then detects and quantifies RNA editing in two passes using JACUSA2 (v2.0.4). The JACUSA2 artefact filters exclude sites adjacent to read starts/ends, INDELs, or splice junctions; sites within homopolymeric runs; and sites with more than two alleles. We run JACUSA2 in single-sample mode for each sample to generate per-base counts at each candidate site.

The first pass, site discovery, identifies putative editing sites in each sample, returning only regions with at least 10x coverage in a given sample. These sites are then converted to VCF format and merged across the cohort. The following filters are applied by default during this first pass: (i) detection in at least 3 samples and in at least 10% of all samples, (ii) a median editing rate of at least 0.01 across detected samples, and (iii) a minimum of 2/3 of samples with detected sites passing JACUSA2’s internal filters. The second quantification pass then performs targeted quantification of RNA editing at the set of discovered sites, ensuring complete per-base counts for all samples. The following stricter filters are applied by default to the sites identified during the second pass: (i) a minimum of 20% of samples with at least 10x coverage per site; (ii) detection in at least 5 samples and at least 20% of all samples; and (iii) a minimum of 2/3 of samples with detected sites passing JACUSA2’s internal filters. The default thresholds may be adjusted as required.

Edi-Set-Flow encodes A-to-I editing sites as either A>G(+) or T>C(−), where the sign represents the strand of origin relative to the reference genome (+ for forward, − for reverse). JACUSA2 is strand-aware and, for stranded RNA-seq, can distinguish A>G(+) and T>C(-) from A>G(-) and T>C(+), the latter of which are not consistent with A-to-I RNA-editing and are more likely to represent genomic SNVs or other artefacts. For unstranded RNA-seq, Edi-Set-Flow instead infers the likely strand, with all A>G sites inferred as A>G(+) and all T>C sites inferred as T>C(-). Edi-Set-Flow also supports APOBEC cytosine-to-uridine RNA editing analyses in principle, with an equivalent strandedness logic applied to these sites (i.e. C>T(+) and G>A(-)); however, we have not tested this use case.

### Edi-Set-Flow: Site annotation

Sites are annotated with genic features using Ensembl VEP^26^ (v114.0) based on the GENCODE annotation source. This annotates each putative editing site with the relevant gene name, transcript identity, affected gene feature (e.g. 5’ UTR, intron, exon, etc.), predicted consequence, and amino acid substitutions in the case of missense variants. Strandedness is also incorporated into the VEP annotation. As this is not a standard feature supported by Ensembl VEP, annotation is performed in three parts: 1) forward-strand annotation, where all putative RNA editing sites assigned to the forward strand (e.g. A>G(+)) are annotated with only forward-strand GENCODE features; 2) reverse-strand annotation, as 1) but for reverse-strand sites (e.g. T>C(-)) and only reverse-strand GENCODE features; and 3) unstranded annotation, where all remaining editing sites without a strand assignment are annotated against all GENCODE features. VEP is run with the “--pick” flag so that the single most relevant gene consequence is selected per editing site. REDIportal^27^ accession numbers and repeat context from the UCSC RepeatMasker files are then added with vcfanno^28^.

### Edi-Set-Flow: Analysis of differential RNA editing

A companion R package, **edisetr**, performs site-wise differential editing analysis with GLMs, using the R function stats::glm. This framework enables modelling the effects of experimental groups, regardless of the number of groups, supporting both simple binary case-control designs and more complex multifactorial designs. We can also include arbitrary fixed-effect covariates to control for confounders such as age, sex, and sequencing batch.

**edisetr** implements three GLM families for site fitting: linear (i.e. Gaussian), binomial, and quasi-binomial. The linear model directly fits the observed editing rate per site. A caveat is that this approach does not account for sequencing depth: samples with lower coverage will have disproportionately high variance and may bias the fit. To address this, we implemented an empirical Bayes shrinkage procedure using the *ebbr* R package. At each site, a beta prior distribution is estimated from the edited and unedited counts across all samples by maximum likelihood estimation, then each sample’s editing rate is shrunk towards this prior according to its coverage, such that low-coverage samples are shrunk to a greater extent than high-coverage samples^29^. The posterior editing rate is then used as the response in the linear model, and the inverse of each sample’s posterior variance as a weight in the linear fit, such that samples with more precise editing-rate estimates (from higher coverage) are weighted more heavily. We refer to this model as “linear-ebayes”, which provides a fourth option for editing rate fitting, in addition to linear, binomial and quasi-binomial.

The binomial and quasi-binomial families are instead fitted directly to the observed edited and unedited base counts per sample, with sample-wise depth an inherent feature of the binomial model. The standard binomial model assumes that variance in observed proportions is fully explained by the binomial probability and the number of trials; however, this assumption is routinely violated in sequencing data, with the excess observed variability between samples known as overdispersion^30^. The quasi-binomial family accounts for this by estimating a dispersion parameter for the model’s residuals and inflating standard errors accordingly, resulting in better-calibrated p-values. This is a lighter-weight modelling approach than the full beta-binomial implemented in REDITs, while still accounting for overdispersion.

Model fitting is performed in parallel across sites using multidplyr, allowing model fitting to scale to tens of thousands of sites when sufficient compute is available. The model fitting results are returned as a series of four tables. The **Summary Table** reports the fitted coefficients, standard errors, test statistics and p-values, giving the direction and magnitude of each covariate effect. The **Margins Table** reports the estimated marginal editing rates and confidence intervals for each experimental group. These are model-based estimates rather than group means, averaged across model covariates. The **Contrasts Table** reports all pairwise between-group differences in editing rate, with standard errors, test statistics and p-values. Marginal estimates and pairwise contrasts are calculated directly from the fitted model using estimated marginal means methodology, implemented in edisetr for computational efficiency.

Finally, the **ANOVA Table** (analysis of variance/deviance) reports the significance of each modelled term and is the primary basis for detecting differential editing between experimental groups. Term-wise statistics in this table are derived from single-term deletion tests comparing the full model with a reduced model excluding each term, using a chi-squared test for the binomial model, and an F-test otherwise (R stats::drop1). For categorical variables with more than two levels (e.g. experimental group), this provides an omnibus test for differential editing across groups. Models are fitted only to experimental groups that meet minimum filtering criteria: by default, each group must have a median coverage of 10x and a median editing rate of 0.001 at the site. Groups that fail these criteria are excluded for the site, and only sites with at least two remaining groups are tested. By default, FDR is controlled using the Benjamini-Hochberg procedure, applied separately for each model term (covariate) across all tested sites (R stats::p.adjust, method = ‘BH’).

### Edi-Set-Flow: Reporting

A summary report of putative editing sites is generated in HTML format using Quarto markdown. This includes a summary of sites that pass or fail customisable filters before model fitting. Quality-control plots show the site-wise and sample-wise median editing rate against median depth, allowing visualisation of coverage-dependent trends and detection of outlier samples. The report also summarises all detected base substitutions (not limited to A-to-I events) including their frequency, editing-rate distribution and repeat context. A-to-I sites are further summarised by predicted gene consequence, and tables list the most highly edited sites within each VEP impact category.

Differential-editing results are summarised using the ANOVA tests for the experimental-group term. Significant sites are visualised in a Manhattan-style plot, with separate panels for sites within *Alu* sequence, other repetitive sequence and non-repetitive contexts. Group-wise estimated marginal editing rates and confidence intervals are plotted alongside observed editing rates for the top differential sites. The report also includes principal-component and clustered heatmap visualisations of editing patterns, together with interactive tables of the top differentially edited sites. In addition, multiQC^31^ is used to aggregate quality control metrics output by mosdepth, RSeQC and fastp in a separate HTML report.

### Analysis of RNA-editing patterns across adult human brain tissues

Unstranded RNA-seq data from adult human brain tissue from the GTEx project were accessed through dbGaP under accession number phs000424.v8.p2. ADAR expression data were derived from GTEx Analysis V10 available at https://gtexportal.org/. These data were used to test and develop the Edi-Set-Flow pipeline. We arbitrarily selected 50 samples from each of nine GTEx brain tissues to evaluate and demonstrate Edi-Set-Flow. The tissues analysed were cerebellum, cerebellar hemisphere, caudate nucleus, nucleus accumbens, putamen, frontal cortex, anterior cingulate cortex, hypothalamus, and hippocampus.

Clustering was performed on estimated marginal mean editing rates for each tissue for significantly differentially edited sites in R as follows. 1) per site, editing rates were converted to median-based Z-scores by subtracting the median marginal editing rate across tissues and dividing by the standard deviation; 2) Euclidean distances between both tissues and sites were calculated; 3) hierarchical clustering was performed using Ward’s method (stats::hclust, method=“ward.D2”). For site-level clustering, we selected four clusters using stats::cutree based on visual inspection of the resulting cluster assignments.

Unless stated otherwise, results presented are based on data processed with the MINIMAL fastp preprocessing profile and the BWA-MEM-2 aligner. Differentially edited sites were classified as biologically relevant if there was a difference in the estimated marginal means across tissues of at least 0.05.

### Validation and benchmarking

A genome-wide ground-truth set of differentially edited sites is not available for benchmarking. Instead, we assessed model calibration using a permutation-based null distribution. This was derived by creating a random permutation of group (tissue) labels once per site, while leaving the model response and covariates unchanged. We deemed a single permutation sufficient because we aggregated results genome-wide across tens of thousands of putative sites. We then combined the permutated and non-permuted results, used the Benjamini-Hochberg procedure to generate q-values, and defined sites with a q-value less than 0.01 as significantly differentially edited. This enabled us to compare model performance using three measures: 1) the number of significant discoveries in the non-permuted data (a proxy for sensitivity); 2) the empirical false discovery rate (FDR), derived from the number of significant sites in the permuted data divided by the number of significant sites in the non-permuted data; and 3) the area under the receiver operating characteristic curve (AUROC). To calculate AUROC, we labelled non-permuted site-level results as positive and permuted results as negative, using Benjamini–Hochberg q-values as the continuous discrimination score. Although this labelling is not based on known ground truth, non-permuted data are expected to be enriched for genuine differential-editing signal, whereas permuting tissue labels removes systematic biological signal and approximates the null distribution. We performed this for a two-way comparison between cortex and cerebellum, and a multi-way comparison across all nine GTEx brain tissues. For the multi-way comparison, we retained sites even when not all tissues met the per-group coverage and editing rate thresholds, so the number of tissues included at each tested site varied depending on the underlying data (Figure S5).

We tested the REDITs beta-binomial-based GLM as implemented in REDIT_regression.R (https://github.com/gxiaolab/REDITs, commit “af65dc8”). We applied REDIT-regression to the same pre-filtered set of sites and data as the comparison edisetr models. REDIT-regression could not be included in the multi-way comparisons because it does not implement an omnibus test for categorical covariates with more than two levels, as in the ANOVA test implemented in edisetr. Instead, it returns a set of k−1 level-versus-reference p-values rather than a single test of the group factor as a whole, so only the two-way comparison could be meaningfully compared.

## Results

### Data preprocessing effects on site discovery

We investigated how data preprocessing and alignment affect downstream detection of RNA-editing sites. For this analysis, we restricted the dataset to 50 GTEx samples from each of the cortex and cerebellum, tissues known to show large differences in RNA editing. Preprocessing with fastp had only a modest effect on the total number of sites detected (Figure S 1). Compared with no preprocessing, STRICT and MINIMAL preprocessing reduced site detection by 6.7% and 0.6%, respectively, with BWA-MEM-2, and by 5.2% and 0.2% with STAR 2-PASS. In contrast, aligner choice had a much larger effect on site yield: with MINIMAL preprocessing, BWA-MEM-2 identified 51% more sites than STAR 2-PASS (57,720 versus 38,273).

We assessed the effects of preprocessing and alignment on differential site detection using a permuted null dataset to estimate the AUROC (Methods). Despite differences in site yield, differential-detection performance was similar across aligners, with median AUROCs of 0.814 for BWA-MEM-2 and 0.808 for STAR 2-PASS. Preprocessing also had little effect on performance, with median AUROCs of 0.808 without preprocessing, 0.812 with MINIMAL preprocessing, and 0.810 with STRICT preprocessing. Based on these results, we have chosen BWA-MEM-2 and MINIMAL preprocessing as the defaults for Edi-Set-Flow.

### Differential models trade sensitivity for FDR control

We evaluated alternative modelling frameworks for differential site detection (Methods). Each test modelled RNA editing at each putative site as a function of tissue, age, and sex. The set of sites detected in the two-way comparison (cortex vs cerebellum) was broadly similar across modelling techniques, with Jaccard similarity indices, defined as the size of the set intersection divided by the size of the set union, ranging from 0.73 (binomial vs linear) to 0.93 (quasi-binomial vs linear-ebayes) (Figure S 4). We found that the binomial model had the highest sensitivity in both comparisons; however, the false discovery rate (FDR) was unacceptably high and not adequately controlled by the Benjamini-Hochberg procedure (Figure 2). This likely reflects the lack of overdispersion correction. The REDIT-regression method also performed poorly in our benchmark, with both the lowest sensitivity and second-highest FDR.

**Figure 2:**
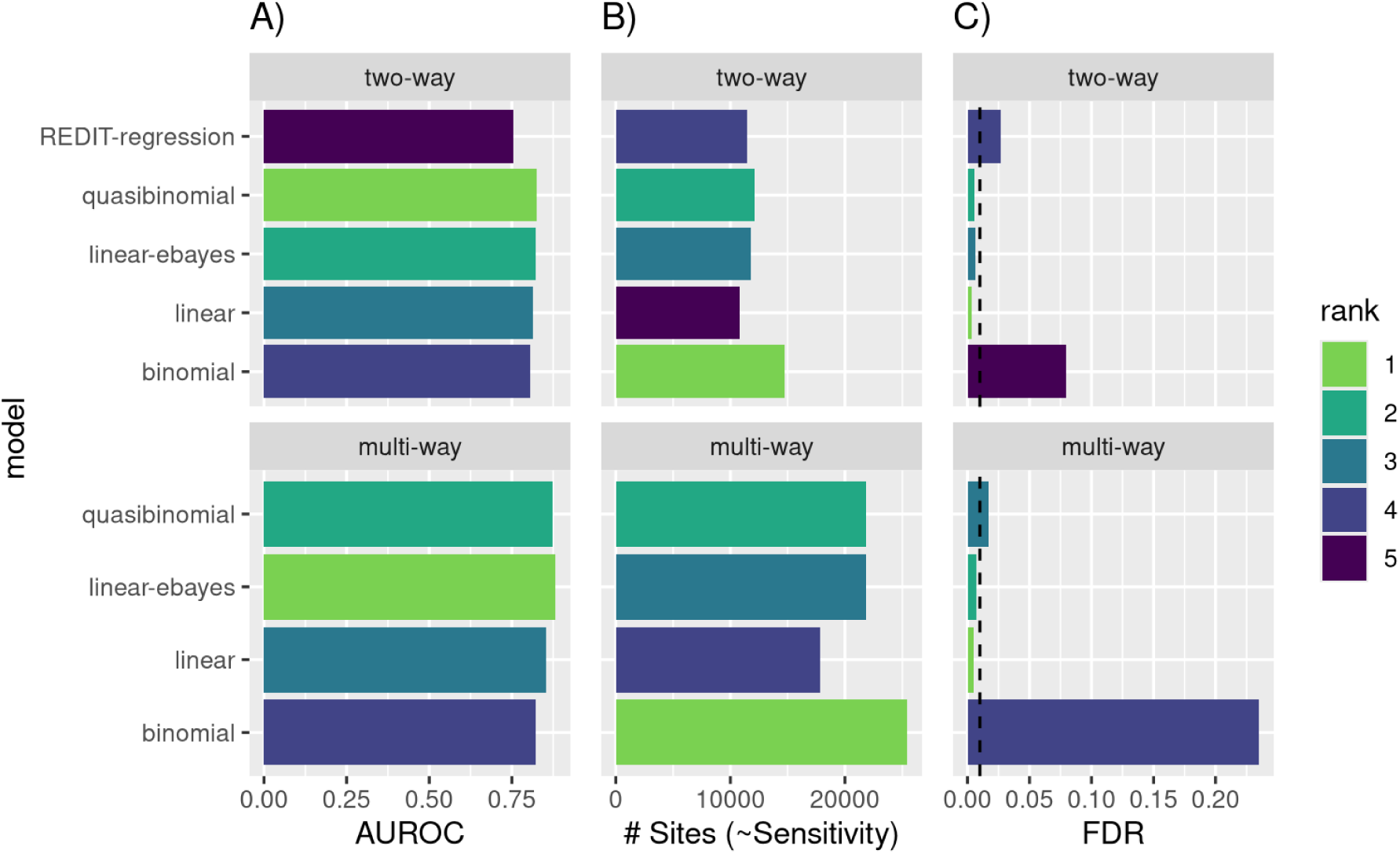
Performance comparison of modelling frameworks for detecting differential RNA editing. Two-way comparisons are based on 50 cortex and 50 cerebellum samples; multi-way comparisons are based on up to nine brain tissues at 50 samples each, with the effective number of groups per site varying due to per-tissue depth and editing rate filtering (Figure S 5). REDIT-regression is shown only for the two-way comparison, as it does not implement an ANOVA-style test for factors with more than two levels. Ranks (best = #1) are indicated on each bar. (A) AUROC for discrimination of permuted-null and non-permuted data. (B) Number of sites detected after applying a Benjamini-Hochberg threshold of q < 0.01. (C) Empirically observed false discovery rate at q < 0.01, estimated from a permutation-based null. The dashed vertical line indicates the nominal FDR (0.01); bars exceeding this line have a higher-than-expected rate of false discoveries.

The quasibinomial, linear, and linear-ebayes models performed similarly well. Empirical Bayes shrinkage boosted the linear model’s sensitivity while maintaining adequate FDR control. Based on the AUROC analysis, the quasibinomial model performed best for the two-way comparison; however, the linear model with shrinkage outperformed it for the multi-way comparison. Analysis of null distribution Q-Q plots confirmed that FDR was inflated for all statistical model families, except the linear and linear-ebayes models (Figure S 6). We prioritised FDR control over the slight sensitivity boost offered by the quasibinomial model; hence, all subsequent analyses are based on the linear model with empirical Bayes shrinkage.

### Edi-Set-Flow detects widespread differential editing across brain tissues

The majority of putative RNA editing sites were A>G or T>C substitutions, consistent with A-to-I editing, and comprised 94% of sites (38,391 / 41,017). These sites had a median editing rate of 0.09, and 82% occurred within *Alu* elements (Figure S 2). The most frequent genic features for A-to-I sites were 3’ UTR (49%), intron (19%) and downstream gene variant (14%) (Figure S3). By default, VEP annotates variants as downstream gene variants when located in the 5 kb immediately downstream of the annotated transcript end (i.e. the end of the 3’UTR).

Statistically significant differential editing across brain tissues was detected for 22,752 of 36,750 candidate sites that had sufficient coverage in at least two tissues (ANOVA Benjamini-Hochberg q-value < 0.01, Figure 3). The median effect size, defined as the maximum difference in editing rate between any two tissues, was relatively low at 0.03. Of these, 7,678 sites (21% of total tested) showed robust differential editing, with an effect size greater than 0.05.

**Figure 3:**
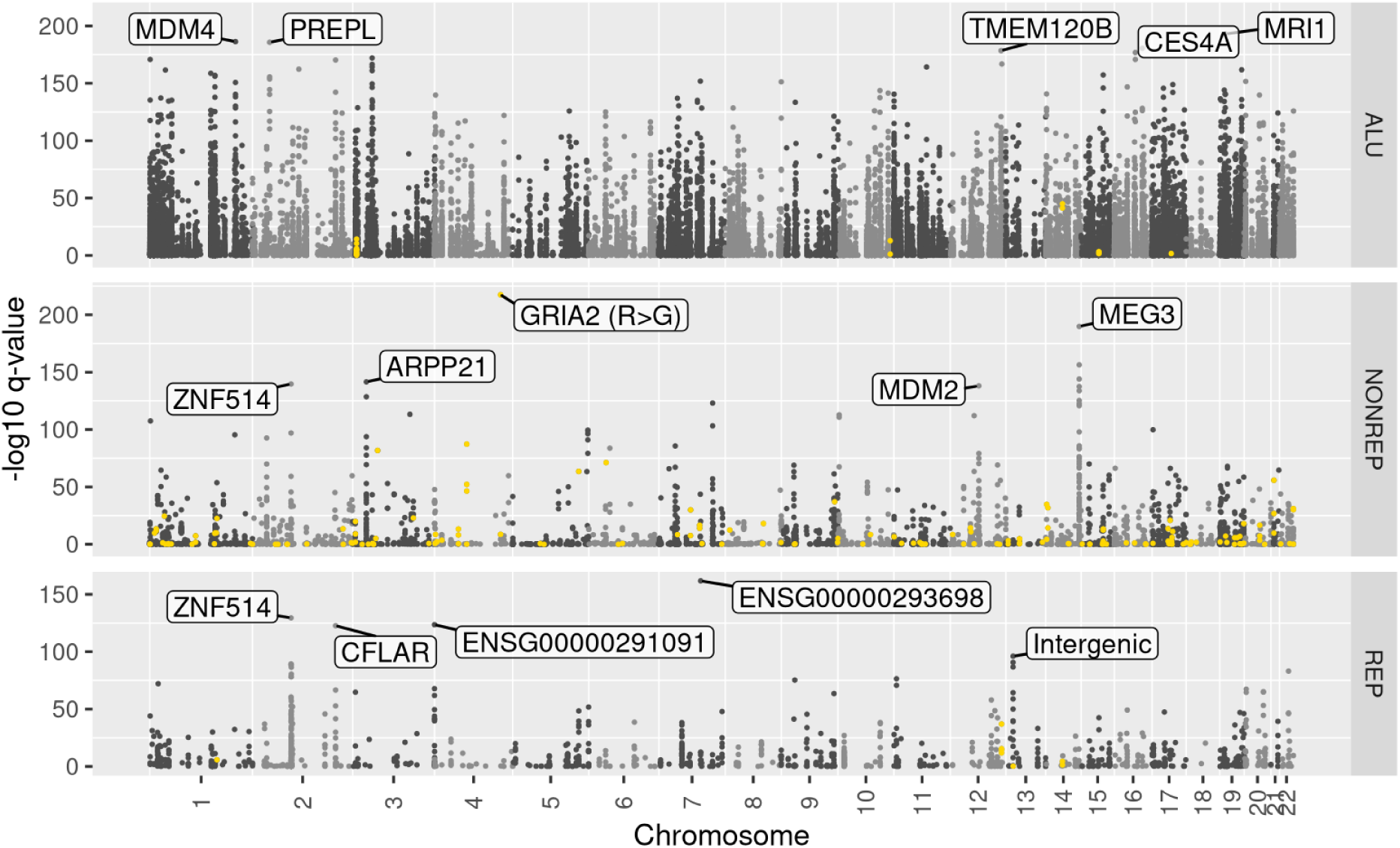
Genome-wide differential editing signatures. Data are faceted by site repeat context; the top 5 genes associated with each sequence context are labelled. Q-values are derived from Benjamini-Hochberg correction of p-values from a linear-ebayes GLM fit ANOVA (F-test). Yellow highlights indicate missense sites. This figure is generated as part of the standard Edi-Set-Flow report.

To identify patterns of differential editing across sites and tissues, we clustered site-wise scaled (z-transformed) editing rates for the 4,454 sites with robust differential editing and sufficient sequencing depth across all tissues. The tissue clustering identified three distinct groupings based on editing patterns: cerebellar (cerebellum and cerebellar hemisphere), striatum (caudate nucleus, nucleus accumbens, and putamen), and the non-striatal forebrain (frontal cortex, anterior cingulate cortex, hypothalamus, and hippocampus) (Figure 4). Site-wise clustering revealed four distinct clusters: clusters 1 and 2 were both characterised by a relative increase in editing in cerebellar tissues, and differed in that cluster 1 sites also displayed a relative decrease in editing in the non-striatal forebrain compared to the striatum. Clusters 3 and 4 were characterised by a relative decrease in editing in the cerebellar tissues, and differed in that cluster 3 sites showed a relative increase in editing in the striatum. These results are broadly consistent with other studies that have analysed GTEx data, with the cerebellum known to have a distinct RNA-editing profile compared to the rest of the brain^6,11^. Significant differences in editing between the striatum and cortex have also been reported at a small number of targeted sites within *HTR2C*, *GRIA2*, *GRIA4*, and *CADPS*^32^.

**Figure 4:**
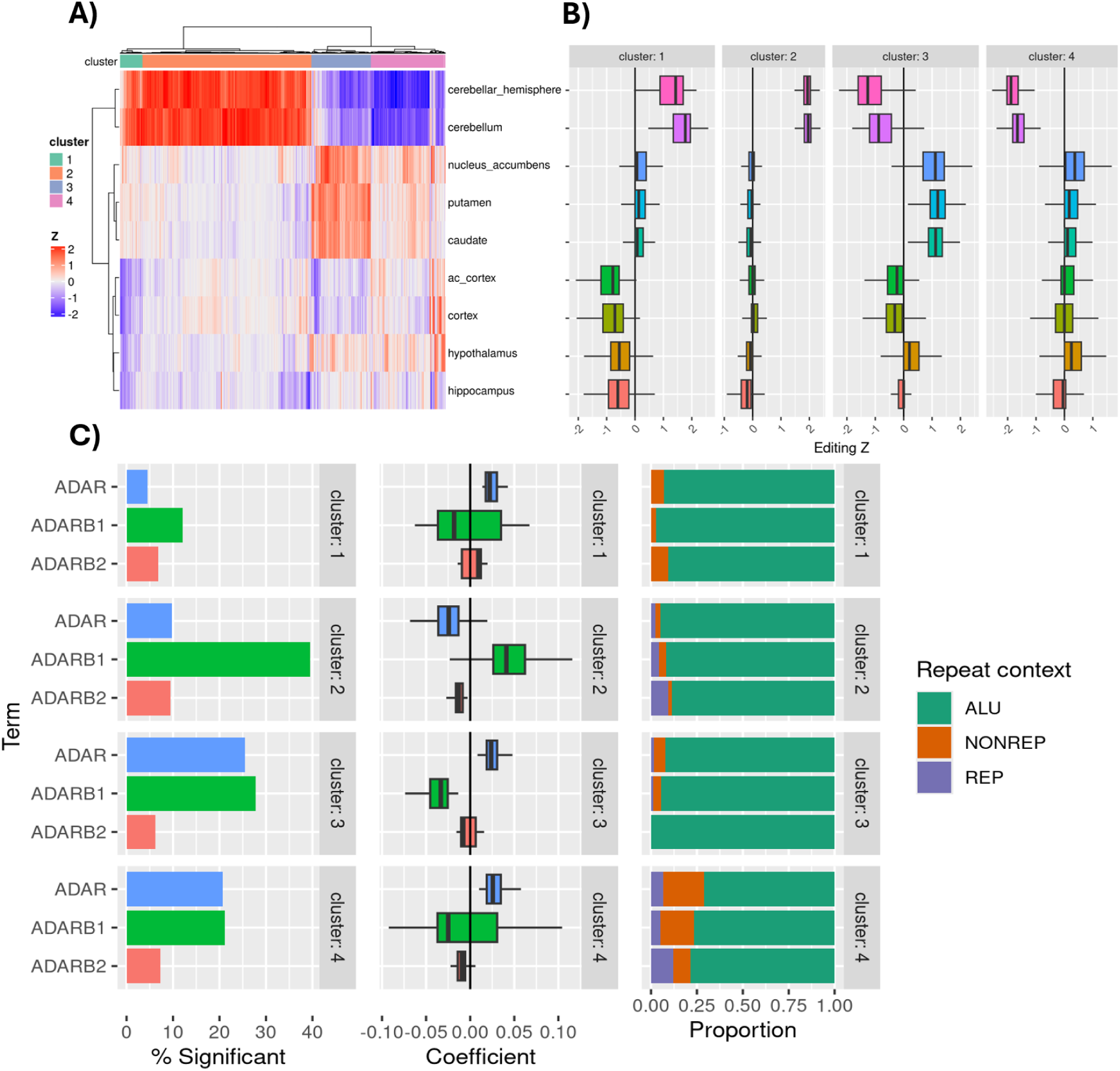
Clustering of differentially edited sites and association with tissue and ADAR expression. A) Heatmap of tissue-wise marginal editing rate z-scores, with clustering based on Euclidean distance using Ward’s method. The x-axis corresponds to individual editing sites, and the y-axis corresponds to tissues. B) Distribution of marginal editing rate z-scores across sites assigned to each cluster in each tissue. C) Significance, magnitude and direction of association of ADAR, ADARB1 and ADARB2 expression with sites in each cluster.

We investigated correlates and potential drivers of editing across these site-wise clusters by fitting nested models at each site: one including only tissue, age, and sex as covariates, and a second adding *ADAR*, *ADARB1*, and *ADARB2* expression covariates (z-scaled log TPM). Cluster 2 sites (increased editing in cerebellar tissues, stable elsewhere) showed the strongest association with *ADARB1*, with approximately 40% of sites nominally associated (p-value < 0.01) with *ADARB1* expression and largely positive association coefficients. Cluster 2 sites also show a negative association with *ADAR* expression, with nominal significance at approximately 10% of sites. The remaining clusters showed a reversal of this trend, with *ADAR* expression positively associated with editing and *ADARB1* expression negatively associated with editing. *ADARB2* appeared to have a smaller effect on editing levels across all clusters, as expected given its catalytic inactivity, with coefficients closer to zero than those of *ADAR* and *<u>ADARB1</u>*.

### Edi-Set-Flow identifies known protein recoding sites

Edi-Set-Flow identified 20 predicted protein recoding events with significant differences in editing rate across the 9 GTEx brain tissues (q-value BH < 0.01) and a magnitude of greater than 0.05 between the highest and lowest tissue editing rates (Table S1). Of these, 18 lie within non-repetitive regions, and two are within *Alu* repeat elements. Both of the recoding sites in annotated *Alu* repeats are in the *SLC38A6* transcript. The magnitude of differential editing was strongly positively correlated with evolutionary constraint at the edited nucleotide, as measured by the 470-way mammalian phyloP conservation score (Spearman ρ = 0.78, p-value = 6.7×10^−5^). This indicates that sites displaying stronger differential editing across tissues are broadly more evolutionarily conserved.

The top five differentially edited sites resulting in protein recoding have been previously described in the literature (Figure 5). The *GRIA2* R764G site showed high editing in cerebellar tissue and the frontal cortex and is associated with *ADARB1* expression. The site causes AMPA receptor channels to both desensitise and resensitize faster, leading to increased sensitivity to glutamate^33^. The *CCNI* R75G site showed lower editing in cerebellar tissues (5%) and higher editing elsewhere (19-22%). *CCNI* encodes a regulator of cell cycle progression^34^. The role of the *CCNI* R75G substitution in the healthy brain is unclear; however, it has been demonstrated to induce the destruction of cancer cells by tumour-infiltrating lymphocytes^35^. The *CADPS* E1250G site shows variability in editing rates across the assayed tissues, with the highest rate in the hypothalamus. Editing at the E1250G site has been shown to increase the release of dense core vesicles from neurons, leading to increased physical activity in mice^36^. The *TMEM63B* Q619R site shows increased editing in cerebellar tissues and is positively associated with *ADARB1* expression. *TMEM63B* encodes an ion channel, and editing has been linked to reduced calcium permeability of the channel^37^. Finally, the *CYFIP2* K320E site showed variability across brain regions, with the cerebellar tissues edited at 71-76%, the cortical tissues at 52-60%, and the remaining tissues at 33-42%, with a strong positive association with *ADARB1* expression. Editing of this site has been shown to play a key developmental role in neuronal axon branching and dendritic spine formation^38^.

**Figure 5:**
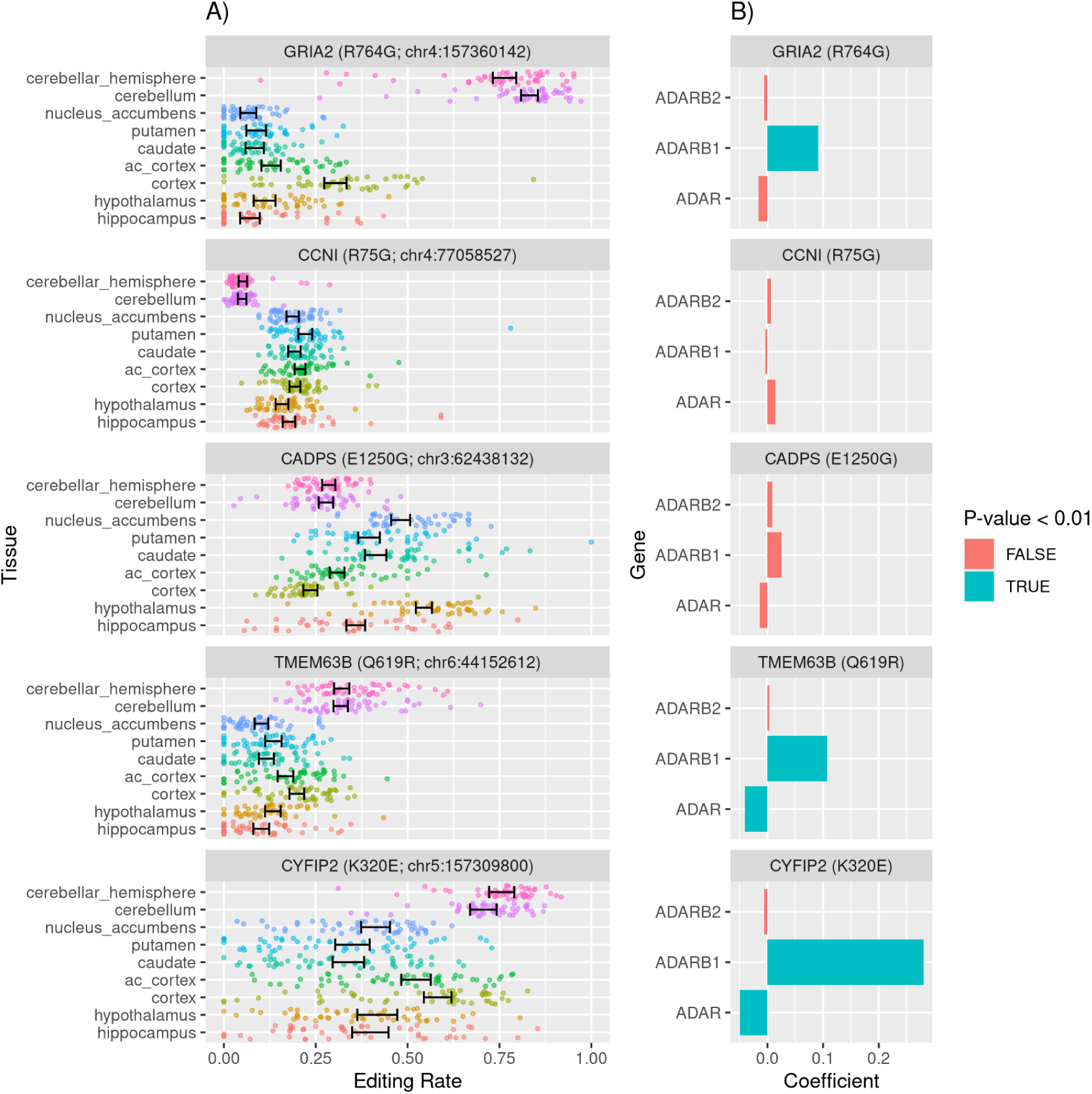
Top differential protein-recoding sites identified across brain tissues. A) Error bars show the modelled marginal upper and lower 95% confidence intervals on tissue-specific editing rates. Jittered points show the underlying per-sample editing rates the model was fitted to. B) Coefficients associated with ADAR, ADARB1 and ADARB2 expression at each site, coloured by whether they obtained nominal significance in the model fit.

## Discussion

We have developed Edi-Set-Flow, a user-friendly end-to-end pipeline capable of robust and sensitive detection of differential RNA editing across multiple experimental groups. Our optimised statistical framework for detecting editing sites, with the option to model and correct for covariates, allows users to robustly detect differentially edited sites while effectively controlling false discovery rates at genome-wide scales. Edi-Set-Flow identifies differential editing across tissues in known protein recoding sites, with the magnitude of differential editing strongly associated with evolutionary conservation.

Our novel approach of applying empirical Bayes shrinkage to raw editing rates before linear modelling proved particularly effective. This was motivated by the observation that although the quasi-binomial GLM improved on the standard binomial model, it still failed to adequately remove the effects of overdispersion in the multi-way comparison, showing residual inflation in the tail of the Q-Q plot that BH correction failed to control, resulting in an elevated false discovery rate (Figure S 6, Figure 2). The use of empirical Bayes was inspired by its effective application to differential gene expression analysis in limma and edgeR^39^. However, unlike these implementations, which pool information across all genes to estimate a shared prior, we estimate the beta prior independently at each site.

Edi-Set-Flow supports both hypothesis generation and hypothesis testing. Hypothesis generation may begin by applying Edi-Set-Flow to RNA-seq data across defined experimental groups and identifying sites with strong patterns of differential RNA editing. For example, observed protein recoding sites can be followed up with experiments that tease apart the functions of edited and unedited protein isoforms in the tissues or samples that show the greatest differential editing. The *CCNI* R75G substitution, observed at lower frequency in cerebellar tissues in the GTEx brain data, currently has no known role in healthy brain tissue but plausibly may fine-tune cell cycle progression, a hypothesis that could be experimentally validated. This edited nucleotide is highly evolutionarily conserved (470 mammal phyloP = 6.1) and is invariant in the human variation database gnomAD v4.1 (n∼800,000).

The tissue-specific editing rate clustering analysis provides further hypotheses that could be explored. The four editing site clusters showed distinct patterns across brain regions, as well as different levels of association with ADAR, ADARB1, and *ADARB2* expression. Knock-in, knock-out, and overexpression experiments in cell lines or organoids could establish whether these associations are causal. For example, sites in cluster 2 appear to be positively associated with *ADARB1* expression and negatively associated with *ADAR* expression, while this trend is reversed for cluster 3. Positive associations are broadly consistent with the established site-specific preferences of ADAR1 and ADAR2^6,11^; however, the basis of the negative associations is less clear and warrants further investigation. In the case of *ADARB2,* negative associations with RNA-editing are well established and are understood to operate through competitive RNA binding at editing sites^40,41^.

In conclusion, Edi-Set-Flow is a statistically robust, scalable and user-friendly tool for accelerating epitranscriptomics research. Edi-Set-Flow is available at <u>GitHub/BahloLab/Edi-Set-Flow</u> under the MIT license.

## Acknowledgements

MB was funded by an Australian National Health and Medical Research Council Investigator grant (APP2033132). BREA was supported by an NHMRC Early Career Fellowship (GNT1157776). JEM was supported by a WEHI-CSL translational data science PhD top-up scholarship. This research was supported by the Commonwealth through an Australian Government Research Training Program Scholarship (DOI: https://doi.org/10.82133/C42F-K220). This work was made possible through the Victorian State Government Operational Infrastructure Support Program and the Australian Government NHMRC IRIISS.

The Genotype-Tissue Expression (GTEx) Project was supported by the Common Fund of the Office of the Director of the National Institutes of Health (commonfund.nih.gov/GTEx). Additional funds were provided by the NCI, NHGRI, NHLBI, NIDA, NIMH, and NINDS. Donors were enrolled at Biospecimen Source Sites funded by NCI\Leidos Biomedical Research, Inc. subcontracts to the National Disease Research Interchange (10XS170), Roswell Park Cancer Institute (10XS171), and Science Care, Inc. (X10S172). The Laboratory, Data Analysis, and Coordinating Center (LDACC) was funded through a contract (HHSN268201000029C) to The Broad Institute, Inc. Biorepository operations were funded through a Leidos Biomedical Research, Inc. subcontract to Van Andel Research Institute (10ST1035). Additional data repository and project management were provided by Leidos Biomedical Research, Inc. (HHSN261200800001E). The Brain Bank was supported supplements to University of Miami grant DA006227. Statistical Methods development grants were made to the University of Geneva (MH090941 & MH101814), the University of Chicago (MH090951, MH090937, MH101825, & MH101820), the University of North Carolina - Chapel Hill (MH090936), North Carolina State University (MH101819), Harvard University (MH090948), Stanford University (MH101782), Washington University (MH101810), and to the University of Pennsylvania (MH101822). The datasets used for the analyses described in this manuscript were obtained from dbGaP at http://www.ncbi.nlm.nih.gov/gap through dbGaP accession number phs000424.v8.p2.

## Supplementary Figures

**Figure S 1:**
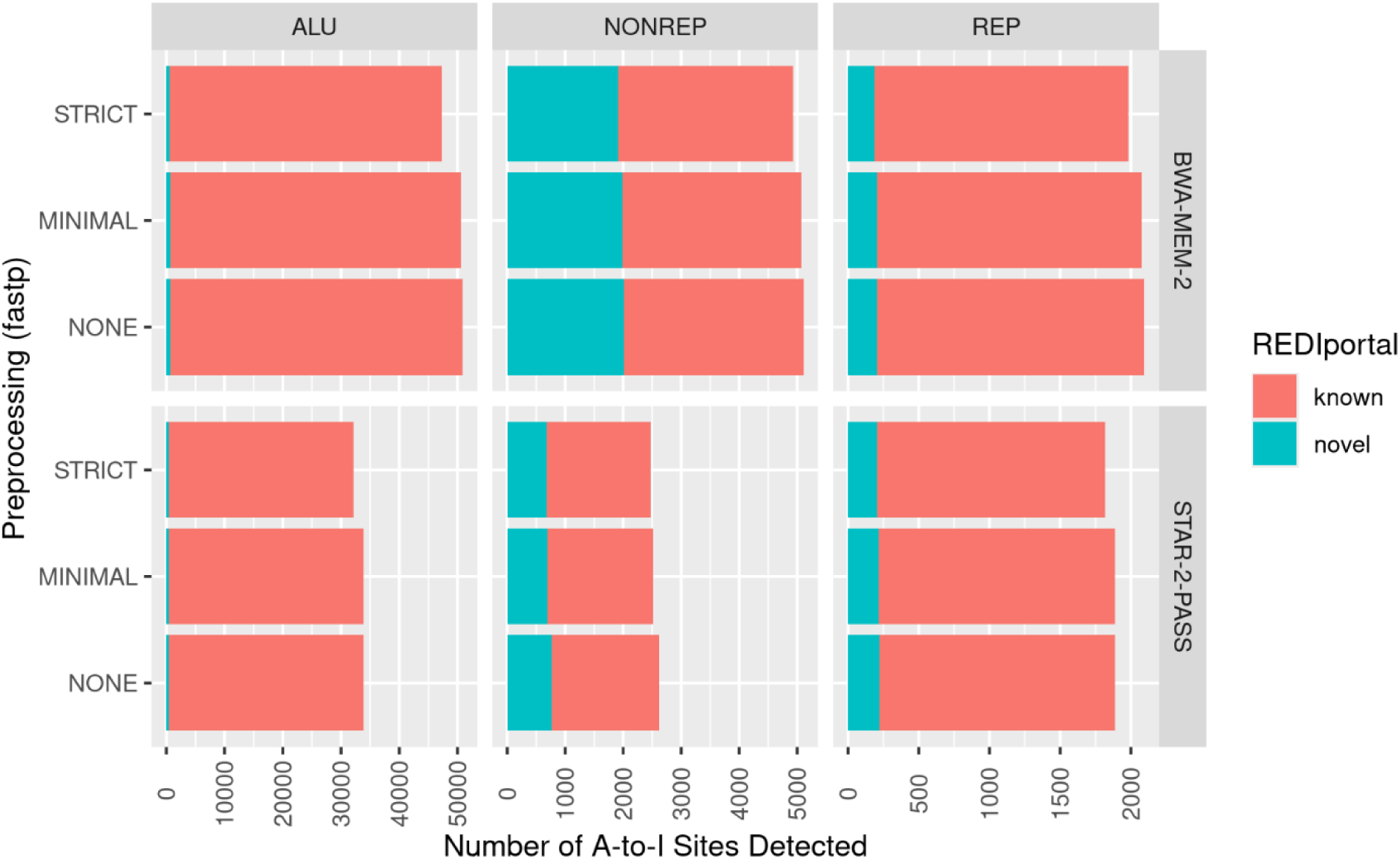
Effects of preprocessing and aligner on site detection. The plots are faceted horizontally by repeat context (ALU – Alu repeat; NONREP – non-repetitive region; REP – repetitive, non-Alu region), and vertically by alignment procedure. Bar plots are coloured based on presence in the REDIportal database. The x-axis of each vertical facet uses a different scale.

**Figure S 2:**
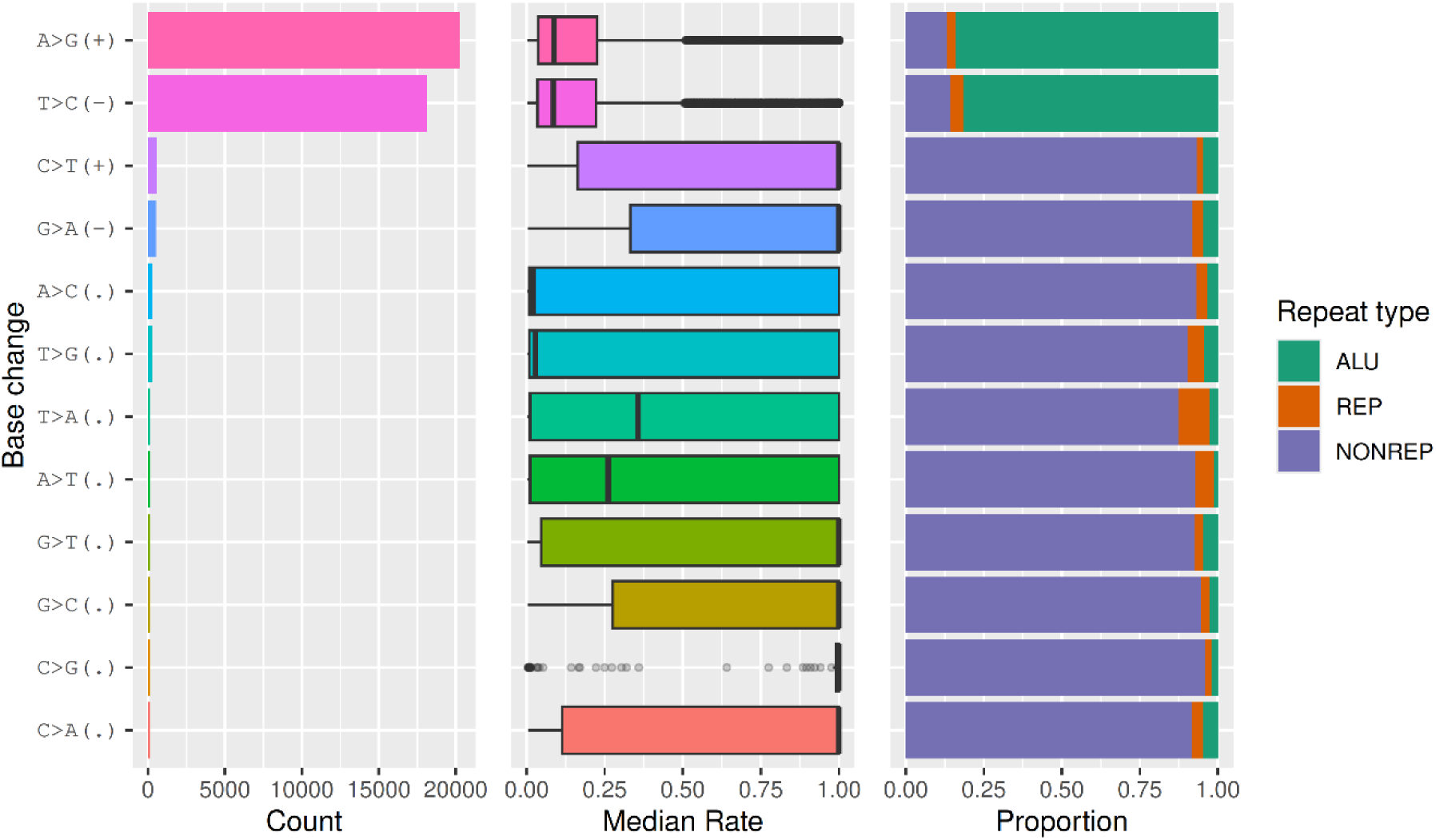
Editing site counts, editing rate distribution, and repeat context by base-change type. This figure is generated by the Edi-Set-Flow report. As the GTEx data is unstranded, base-change strand is inferred only for sites matching the ADAR A-to-I signature (A>G(+) and T>C(-)) and the APOBEC C-to-U signature (C>T(+), G>A(-)).

**Figure S 3:**
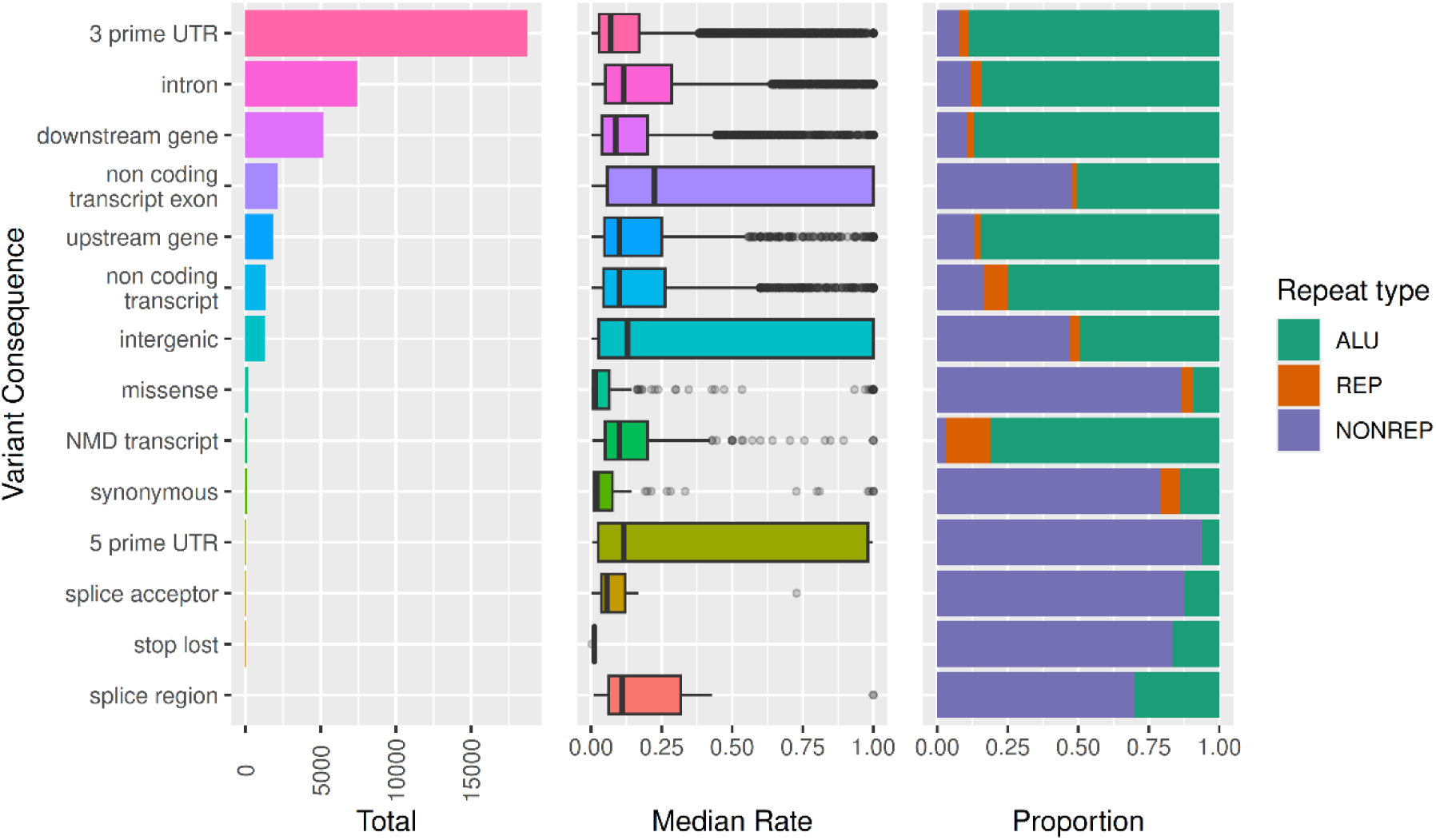
A-to-I editing site counts, editing rate distribution, and repeat context by variant consequence. This figure is generated by the Edi-Set-Flow report.

**Figure S 4:**
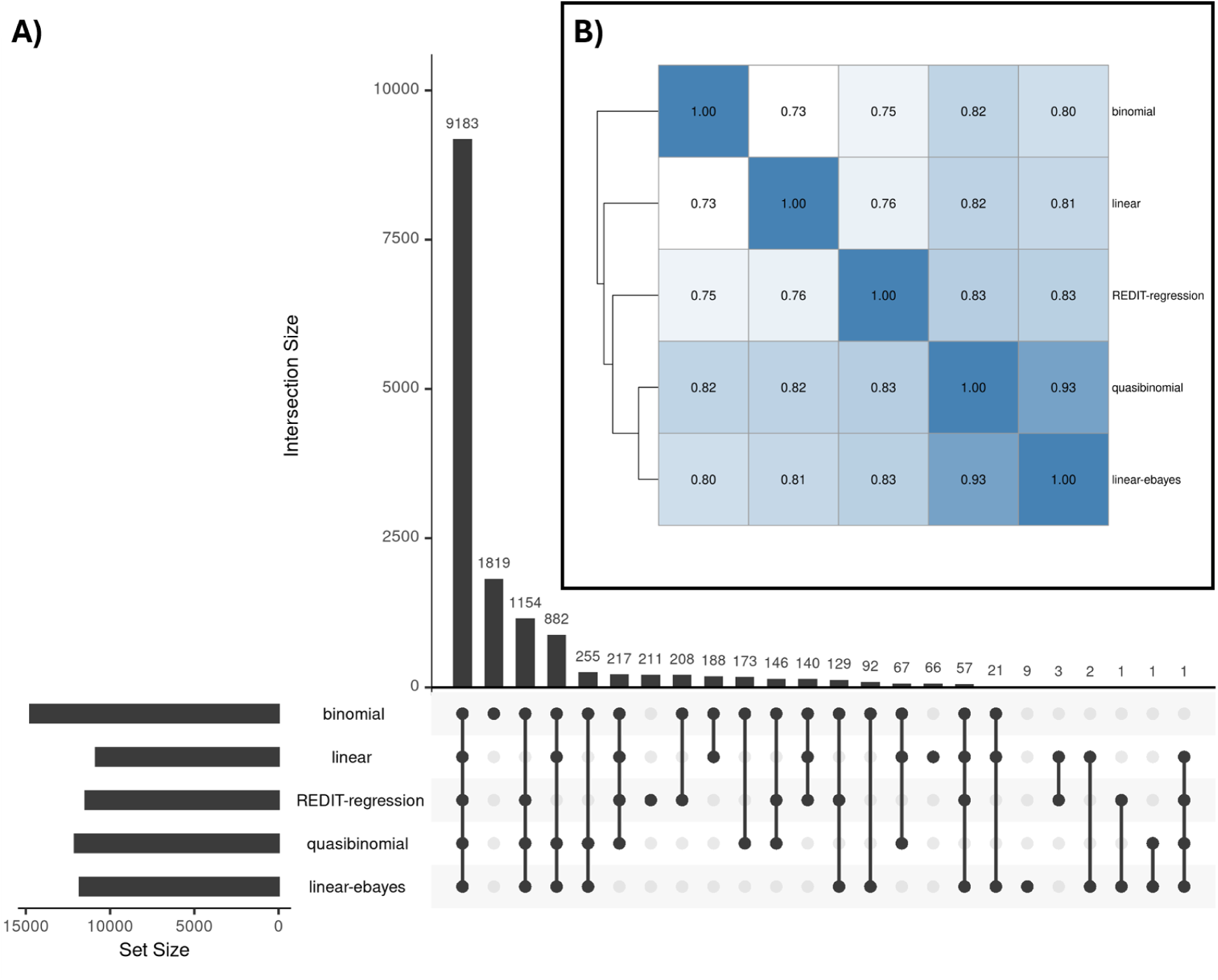
Comparison of differentially edited site sets detected between frontal cortex and cerebellum with different modelling frameworks. Differentially edited site sets defined by Benjamini-Hochberg q-value < 0.01. A) Upset plot, showing the counts identified by each model and each intersection of models. B) UPGMA clustered heatmap of the pairwise Jaccard similarity between each set of sites.

**Figure S 5:**
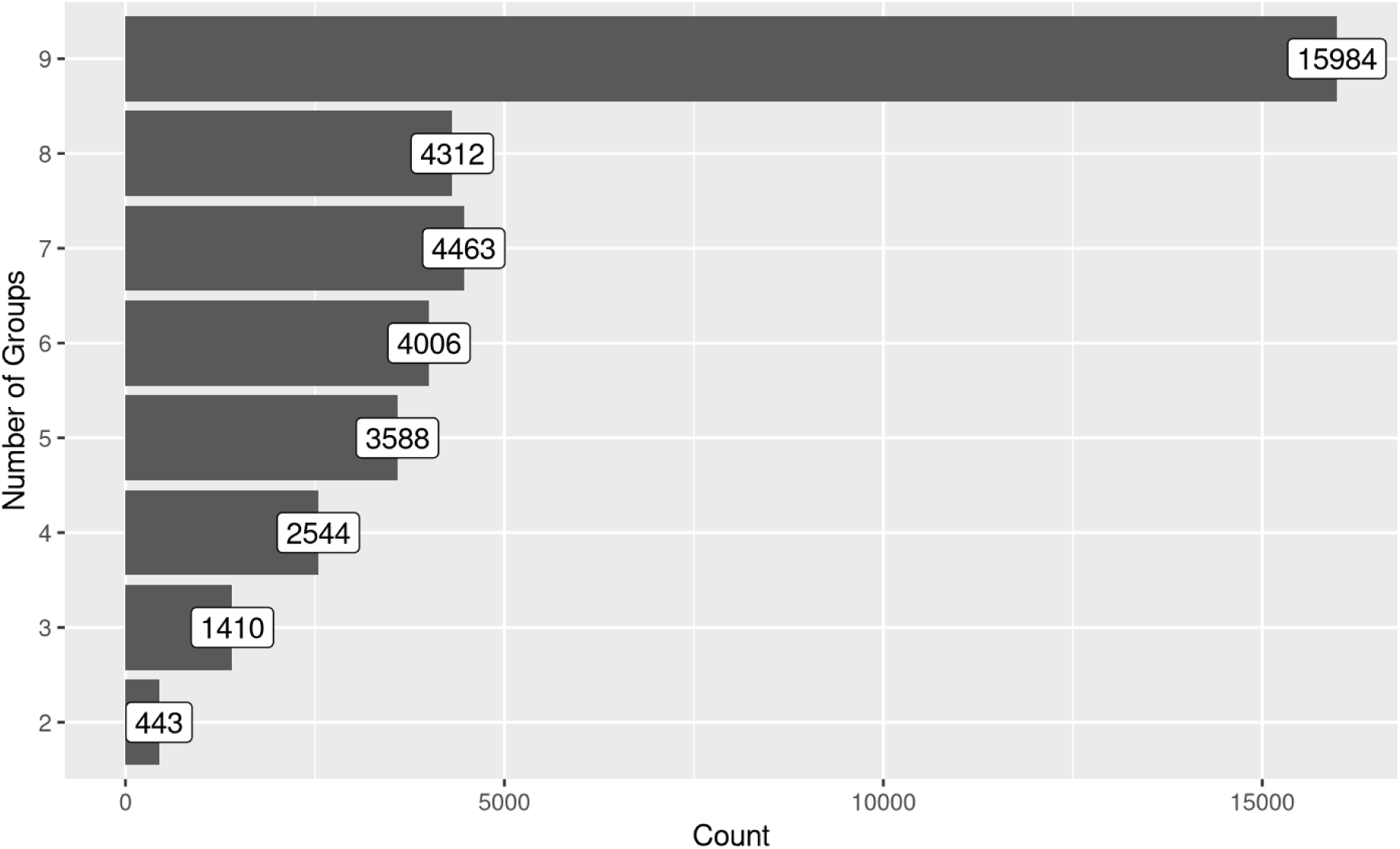
Number of groups after filtering was applied to each editing site and group for the multi-way ANOVA tests. The majority of sites had sufficient coverage for all 9 GTEx brain tissues to be included in the differential tests.

**Figure S 6:**
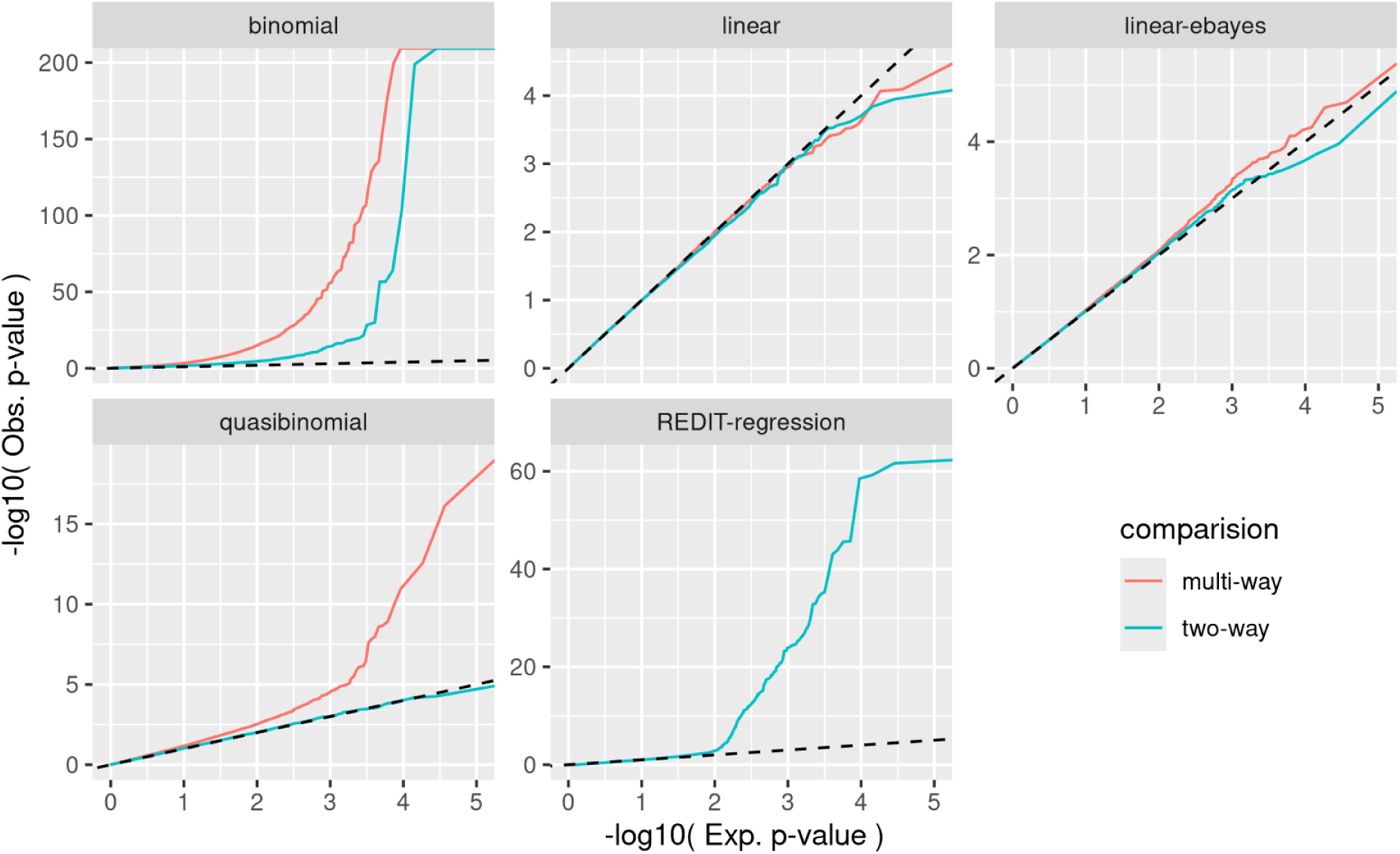
P-value Q-Q plot based on the permuted null distributions using different modelling frameworks. The x-axis scale is consistent across facets; however, the y-axis scale varies by dataset. The dashed line is along the function y=x. Elevation of points above this line indicates that p-values are inflated compared to the expectation under the null hypothesis.

